# *Deformed wings* is an Atg8a-interacting protein that negatively regulates autophagy

**DOI:** 10.1101/2023.02.03.526972

**Authors:** Marta Kołodziej, Panagiotis Tsapras, Aristi Konstantinou, Alexander D. Cameron, Vasilis Promponas, Ioannis P. Nezis

## Abstract

LC3 (microtubule-associated protein 1 light chain 3, called Atg8 in yeast and *Drosophila*), is one of the most well-studied autophagy-related proteins. LC3 controls selectivity of autophagic degradation by interacting with LIR (LC3-interacting region) motifs also known as AIM (Atg8-interacting motifs) on selective autophagy receptors that carry cargo for degradation. Non-degradative roles of LIR motif-dependent interactions of LC3 are poorly understood. Here, we used yeast-two hybrid screening and Alpha Fold structural predictions, and we identified transcription factor Dwg (Deformed wings) as an Atg8a-interacting protein in *Drosophila*. Dwg-Atg8a interaction is LIR-motif dependent. We further show that Dwg is a negative regulator of autophagy. Our results provide novel insights on non-degradative roles of LIR motif dependent interactions of Atg8a and the transcriptional regulation of autophagy in *Drosophila*.

## Introduction

Autophagy is an evolutionarily conserved lysosomal degradation process involved in breakdown and recycling of intracellular components (Lamb et al., 2013; Feng et al., 2014). There are three main types of autophagy: macroautophagy, microautophagy and chaperone-mediated autophagy; all of which rely on the lysosome to digest intercellular cargo (Parzych & Klionsky, 2014). During macroautophagy, a double-membrane phagophore is formed to surround and isolate cargo. The enclosed cargo is then transported to the lysosome where it is broken down by hydrolytic enzymes and its components are returned to the cytosol to be re-utilized by the cell (Lamb et al., 2013; Feng et al., 2014). Macroautophagy is the most studied type of autophagy and will hereafter be referred to as simply autophagy. Upregulation in the autophagy pathway is typically a survival response to nutrient depravation, although autophagy is constantly active at basal levels in most cell types allowing for general housekeeping functions and protection of intracellular integrity (Lamb et al., 2013). Autophagy also plays an important role in quality control of proteins and organelles, making it a crucial process in maintenance of homeostasis, dysfunction of which can lead to abnormal protein build up and many pathologies (Yamamoto et al., 2023).

Autophagy was considered to be regulated exclusively by cytosolic processes, however increasing evidence over the years has shown that transcriptional and epigenetic events are crucial to autophagy regulation. Although not much is currently known about transcriptional regulation of autophagy-related (ATG) genes, several transcription factors have been suggested to activate or suppress expression of *Atg* genes in *Drosophila* and yeast (He & Klionsky, 2009; Shu et al., 2020). Following the discovery that transcription factor EB (TFEB) can regulate a wide range of autophagy-related genes in response to nutrient deprivation, research into transcriptional regulation of autophagy has significantly increased, leading to the discovery of many additional transcription factors directly involved in autophagy regulation. Discovery of transcription factors that enhance the expression of autophagy genes also allow for the possibility that autophagic activity can be regulated from the nucleus (Shu et al., 2020; Ma et al., 2022). Transcriptional dysregulation in turn is often the cause of many human syndromes and complex diseases, including autoimmunity, neurological conditions, and developmental disorders, frequently due to signalling pathways targeting transcription machinery linked to autophagy regulation (Shu et al., 2020).

Microtubule-associated protein 1 light chain 3 (LC3; known as Atg8 in *Drosophila*) is one of the most studied autophagy related proteins. Among other ATG proteins, LC3 is essential in autophagy and is required for elongation and maturation of the autophagosome. In recent years the LC3-interacting region (LIR) or Atg8-interacting motif (AIM) has also been discovered as an autophagy receptor, providing invaluable insight into the function of selective autophagy (Pankiv et al., 2007). Selective autophagy receptors (SARs) allow cargo to be degraded by interacting with Atg8 family proteins like LC3 via short amino acid LIR sequences, which bind to the Atg8 proteins by the LIR docking site (LDS) inducing autophagosome formation. Many ATG8/LC3 interacting proteins contain a hydrophobic LIR motif with a core sequence of (W/F/Y) XX(L/I/V) (Adriaenssens et al., 2022). Experimental validation of several LIR motifs triggered the development of *in silico* tools to predict new LIR motif instances such as iLIR (Kalvari et al., 2014), hfAIM (Xie et al., 2016) and pLIRm (Han et al., 2021). Although these computational tools have been successfully used to predict LIR motifs, structural determination of the spatial distribution of LIR motifs or flanking residues on ATG8 proteins is missing.

The protein structure prediction tools AlphaFold2 (AF2) (Jumper et al., 2021), AlphaFold-Multimer (AF2-multimer) (Evans et al., 2022) and Uni-Fold (Li et al., 2022) have revolutionized the use of structural biology in understanding complex molecular interactions. In this study, we used AlphaFold and Uni-Fold to examine if they could accurately predict ATG8-binding structures mediated by LIR motifs in experimentally verified LIR motif-containing proteins (LIRCPs) in *Drosophila melanogaster*. Both AF2 and Uni-Fold demonstrated high accuracy in determining LIR motifs in experimentally validated proteins that carry functional LIR motifs in *Drosophila*. By utilizing high-throughput yeast-two-hybrid (Y2H) screen and the predictive accuracy of AF2/Uni-Fold, we show here that the transcription factor *deformed wings* (Dwg) is a novel Atg8a-interacting protein that binds Atg8a in a LIR motif-dependent manner. We also demonstrate that Dwg functions as a transcriptional suppressor of autophagy and that depletion of Dwg induces autophagy in nutrient-rich conditions through the enhanced expression of autophagy genes.

## Results and Discussion

### Prediction of experimentally validated UR motifs using AF2/UniFold

GABARAP is the closest mammalian homolog of the Dmel Atg8a (Identity: 107/117 [91.5%]). We verified that AlphaFold reasonably models GABARAP. For this, we compared the AF model available from the UniProt entry (AF-O95166-F1-model_v3) and compared it to the experimentally determined structure (1GNU) using TM-align (Zhang & Skolnick, 2005). The resulting structural alignment (TM-score = 0.96) indicates a near perfect match (Figure 1A). Similar results were obtained when the AF model was compared with the GABARAP structure bound to the ULK1 LIR-motif (6HYO; TM-score=0.94).

**Figure 1.**
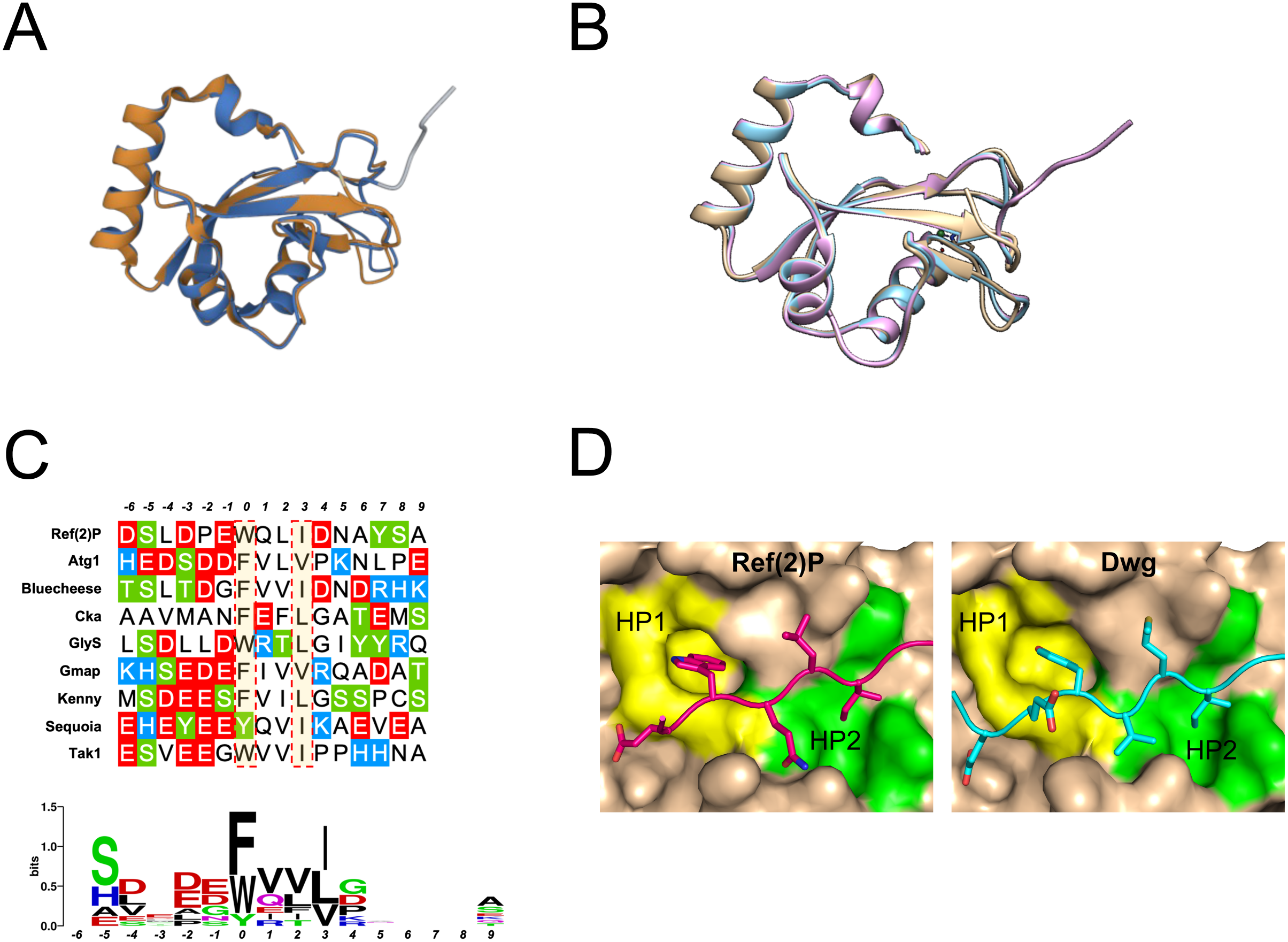
Structural characterization of LIR/LDS interactions using Alpha Fold/Uni-Fold. A) Superimposition of AF model for *Drosophila* Atg8a (brown; AlphaFold: AF-Q9W2S2-F1-model_v3) model with experimentally determined human GABARAP structure (blue; PDB: 1GNU). B) Superimposition of the UniFold models for *Drosophila* Atg8a (purple) and human GABARAP (blue) against the experimentally determined human GABARAP structure (brown; PDB: 1GNU). C) LIR motif consensus of known *Drosophila* LIRCPs. (Top panel) Alignment of the experimentally verified functional LIR motifs identified in *Drosophila* LIRCPs. The LIR peptide extends 6 amino acids N-term. and 9 amino acids C-term. from the Atg8a LDS HP1-binding residue (designated as position “0”). Highlighted red dotted boxes denote the critical amino acids at positions “X0” and “X3” of the LIR core that dock to the HP1 and HP2 sites respectively, within the Atg8a LDS pocket. Acidic amino acids are represented with red, basic amino acids with blue, and with green are Ser and Thr residues than may be targets of de/phosphorylation cycles. (Bottom panel) Sequence logo map for the conservation of each residue with respect to its position in the extended LIR motif. Sequence logo graph constructed using WebLogo (available at: https://weblogo.berkeley.edu/). D) Superimposition of Ref(2)P (left) and Dwg (right) protein backbones docking to the HP1 (yellow) and HP2 (green) sites of Atg8a.

Due to the fact that AlphaFold-multimer predictions are quite slow along with limitations on the length of sequences that can be analyzed, we used Uni-Fold instead (Li et al., 2022). For this, we compared the best Uni-Fold model (based on pLDDT score) obtained using https://colab.research.google.com/github/dptech-corp/Uni-Fold/blob/main/notebooks/unifold.ipynb (Mirdita et al., 2022)

We performed the comparison of the Uni-Fold model for the *Drosophila* Atg8a to the human GABARAP experimentally determined structure (1GNU). The resulting structural alignment (TM-Score=0.96) again showed a high fit between the predicted model and the experimental structure (Figure 1B). Inspection of the superposed structural models of Atg8a (purple) and GABARAP (blue) against the experimental GABARAP structure (1GNU, brown) illustrated that the regular secondary structural elements are very well conserved, with any differences residing in loop regions (Figure 1B). In addition, residues within the two hydrophobic pockets (HP1: E17, L50, F104; HP2: V51, L55, L63) which are known to be important for LIRs binding to GABARAP (Rogov et al., 2017), are conserved when a structure-imposed sequence alignment of Atg8a and GABARAP is considered.

To examine the structural details of LIR motif/Atg8a interactions in *Drosophila, we* run Uni-Fold models using experimentally validated, canonical (W/Y/ F-X-X-L/I/V) LIR motifs (Figure 1C). The Uni-Fold-predicted models showed high accuracy of experimentally verified LIR motifs in most of the proteins (Table 1 and Figure 1D). Taken together the above data highlight that AF/Uni-Fold are valuable tools to assist the prediction and characterization of functional LIR motifs.

### Dwg is an Atg8a-interacting protein and transcriptional repressor of autophagy

To identify novel Atg8a-interacting proteins in *Drosophila*, we performed a Y2H screening using *Drosophila* Atg8a (1-121), as a LexA-bait (pB27) and an inducible LexA-bait fusion (pB31), performed on a *Drosophila* 3^rd^ instar larvae library. One of the positive hits is the transcription factor Dwg (also known as Zw5) (Figure 2A). We used Uni-Fold to explore the interaction between Atg8a and Dwg. This showed that Y129A and I132 docked into the HP1 and HP2 sites of Atg8a respectively, similar to what is observed for Ref(2)P (Figure 1D).

**Figure 2.**
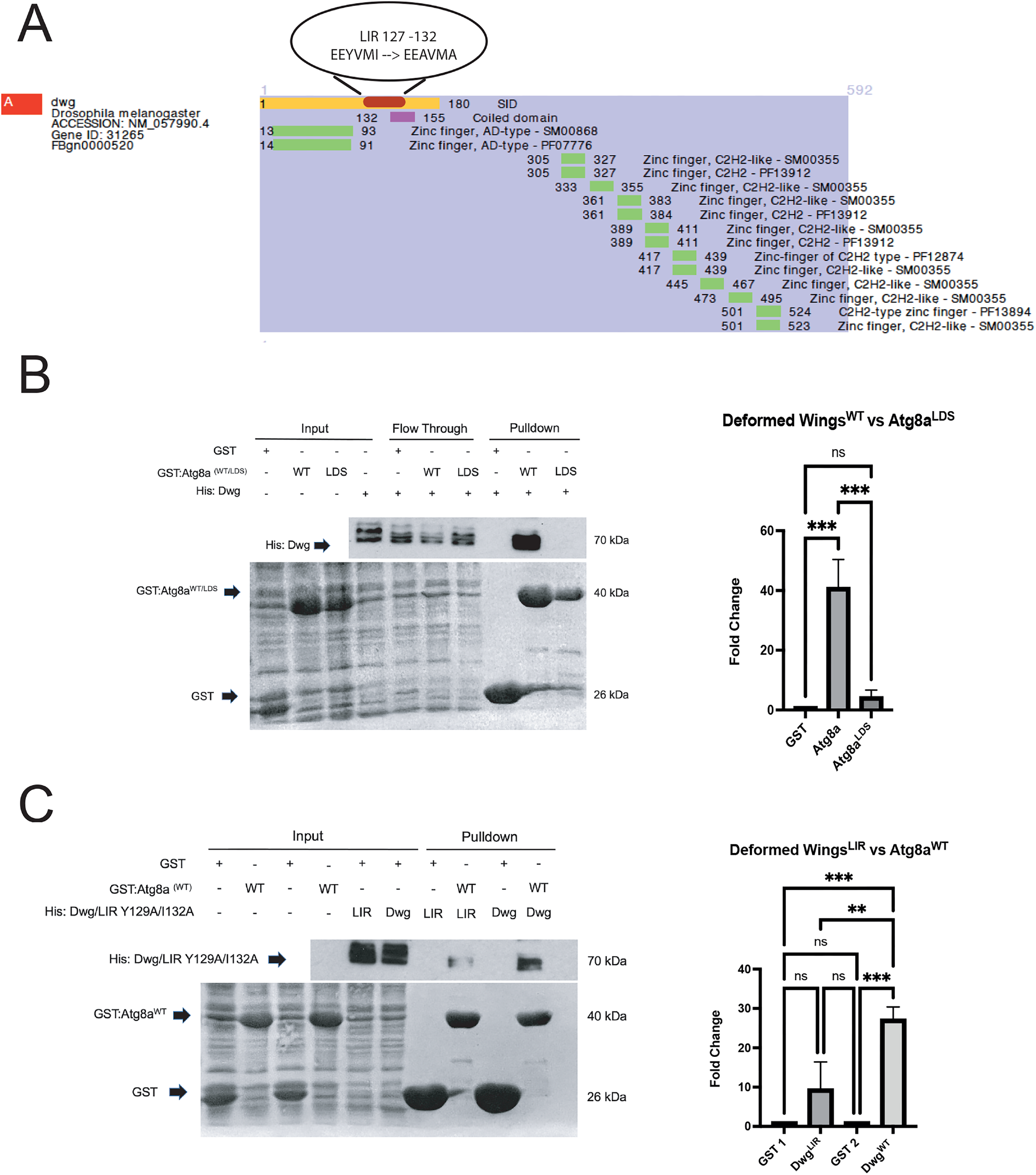
Dwg is an Atg8a-interacting protein. A) Yeast-two-hybrid data showing interaction of the Selected Interaction Domain fragment (SID) in yellow in the region between 1-180 of Dwg with Atg8a. Using the online iLIR software, a LIR motif with the sequence EEYVMI was identified within the SID region at position 127-132. B) GST-Pulldown assay confirming interaction between Dwg and Atg8a^WT^ and showing a reduced interaction between Dwg and Atg8a^LDS^ mutant. The LDS contains a mutation in the docking site on Atg8a thereby preventing LIR dependent interactions. Experiment was performed in triplicate and data was analysed by one-way ANOVA with a post hoc Tukey test, ***p< 0.001, ns: not significant based on the statistical analysis performed. C) GST-pulldown assay looking at the interaction of Dwg with Atg8a when a mutation has been introduced in the predicted LIR motif responsible for the Atg8a binding. Alanine substitutions were introduced in the predicted binding sequence at positions 129 and 132 (EEYVMI → EEAVMA). Experiment was performed in triplicate and data was analysed by one-way ANOVA with a post hoc Tukey test, ***p< 0.001, **p< 0.01, ns: not significant based on the statistical analysis performed.

We confirmed the direct interaction between Dwg and Atg8a using glutathione S-transferase (GST)-pull-down binding assays (Figure 2B). This interaction was significantly reduced when we used a mutant of Atg8a in which the LIR motif docking site (LDS) (Y49A) was impaired, indicating that the interaction between Dwg and Atg8a is LIR motif dependent Furthermore, point mutations of the Dwg LIR motif in positions 129 and 132 by alanine substitutions of the aromatic and hydrophobic residues (Y129A and I132A) reduced its binding to Atg8a (Figures 2B and 2C). These results, show that the LIR motif at position 129-132 mediates the interaction between Dwg and Atg8a. Given the observed interaction between Dwg and Atg8a, we examined whether Dwg colocalizes with Atg8a. Immunofluorescence analysis showed that Dwg colocalizes with Atg8a in the nucleus upon fed conditions and in cytoplasmic puncta under starved conditions (Figure 3A and 3B). These results indicate that Dwg is an Atg8a-interacting protein, and that this interaction is LIR motif-dependent.

**Figure 3.**
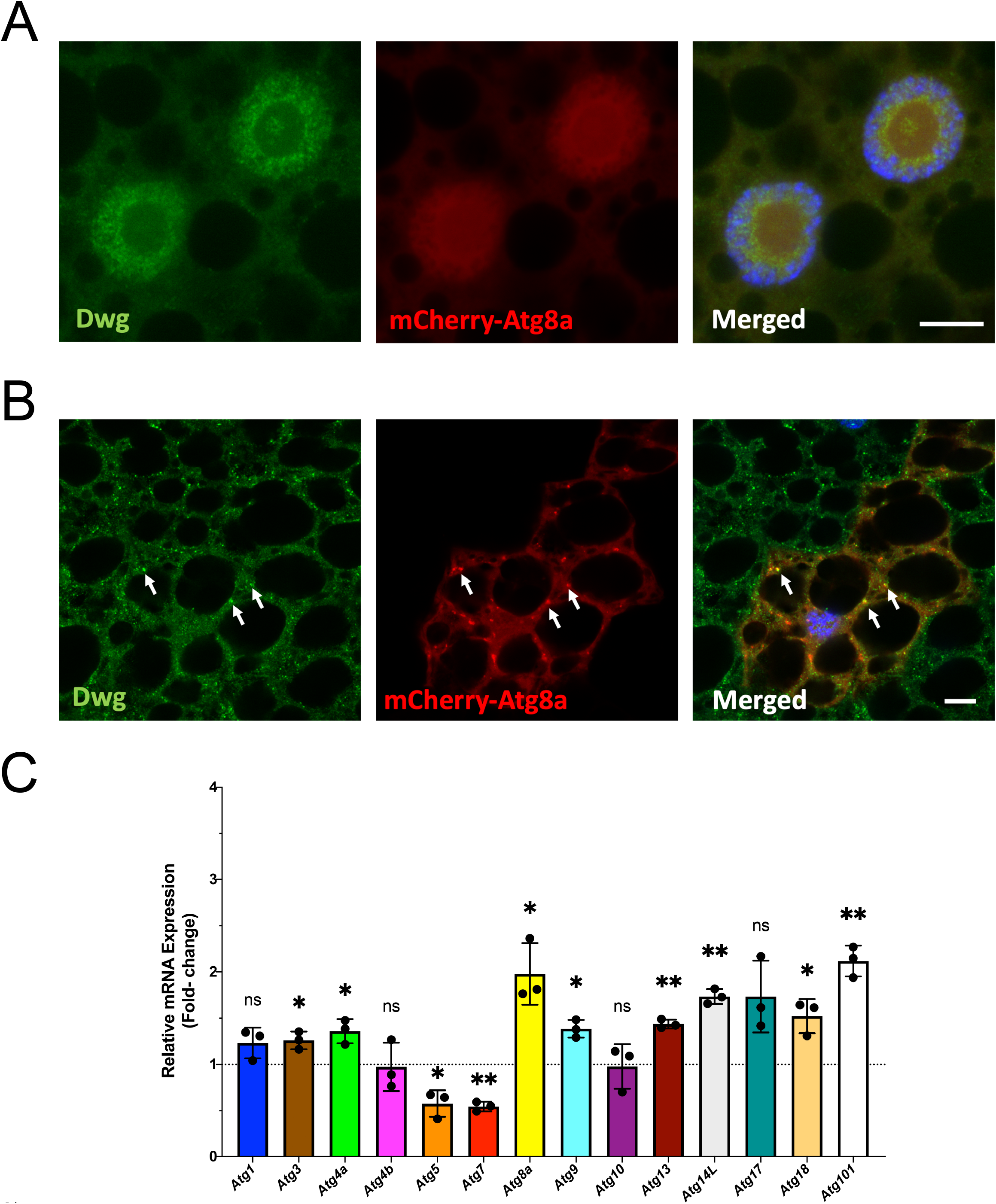
Dwg is a negative regulator of autophagy. A) Confocal micrographs of larval fat bodies under fed conditions expressing mCherry-Atg8a and stained for endogenous Dwg (green). Scale bar 10 microns. B) Confocal micrographs of larval fat bodies under starved conditions (4 hours, 20% sucrose) expressing mCherry-Atg8a and stained for endogenous Dwg (green). Arrows show colocalization between cytoplasmic Dwg puncta and the autophagic marker mCherry-Atg8a. Scale bar 10 microns. C) Expression of key *Atg* genes in *Dwg^8^* mutants, measured by qPCR. The Ct of each gene was measured as average from triplicate wells and graph constructed using the ΔΔCt method. 1) Values >1 depict upregulation, while values <1 or =1 indicate downregulation or no change in expression respectively, relative to the female control group. Data shown as mean + SD (n =3 individual biological repeat; ns, not significant; *, p<0.05; **, p<0.01).

We next examined whether the expression of autophagy genes was affected in Dwg mutants. Real-time quantitative PCR (qPCR) analysis showed that the expression of numerous core autophagy genes was increased (Figure 3C) suggesting that Dwg is a transcriptional repressor of autophagy. Our results demonstrated that *Atg8a* exhibited the highest fold-change in upregulation in Dwg mutants, followed by *Atg101* and *Atg14* (Figure 3C). Other *Atgs* important for autophagy initiation, Atg8a lipidation and autophagosome formation, such as *Atg3, Atg4a* and *Atg9* (Lamb et al., 2013), respectively, also displayed significant upregulation among others (Figure 3C). Our results are consistent with recent findings shown that Dwg is transcriptional repressor of autophagy (Tang et al., 2022). Taken together, our qPCR-based observations suggest that Dwg suppresses transcriptional expression of core autophagy genes in fruit flies under normal conditions.

In this study we highlight a non-degradative role of LIR motif-dependent interactions of Atg8a in Drosophila. We also show that Alpha Fold simulations are very useful for studying LIR-LDS interactions. Finally, we have uncovered that transcription of autophagy genes can be regulated by the interaction of transcription factor Dwg with Atg8a. Our data provide important information on the transcriptional regulation of autophagy in *Drosophila*.

## Materials and Methods

### Analysis using AF/Uni-Fold

Sequences of proteins analysed were obtained from UniProt. Superposition of AF/Uni-Fold predictions on known structures was performed using the align command in PyMOL (The PyMOL Molecular Graphics System, Version 2.3.5 Schrödinger, LLC).

### Fly husbandry and generation of transgenic lines

Flies used in experiments were kept at 25°C, 70% humidity and raised on a cornmeal-based diet. The *dwg^8^* (BDSC4094) flies were obtained from Bloomington *Drosophila* stock centre. The mCherry-Atg8a flies have been described previously (Nezis et al., 2009). Yeast-two-hybrid screening was performed as described in Tsapras et al., 2022.

### Immunohistochemistry

Larva fat bodies were dissected in PBS and fixed for 30 min in 4% formaldehyde at room temperature. Blocking, as well as primary/secondary antibody incubations were performed in PBT (0.3% BSA • 0.3% Triton-X100 in PBS). Primary and secondary antibodies were incubated overnight at 4 °C, or for 2 hrs at room temperature, in PBT.

Anti-Dwg (anti-Zw5, DSHB) primary was used at 1:5 dilution. Hoechst 33342 DNA staining dye (New England Bioloabs #4082, 1:1000 in PBS) was used to visualize nuclei. Washes were performed in PBW (0.1 % Tween-20 in PBS).

All images were acquired in Carl Zeiss LSM710 or LSM880 confocal microscopes, using a 63x Apochromat objective.

### Protein expression and purification for GST pulldown assay

Both, the GST-fusion bait, and the His-labelled prey proteins were expressed in *Escherichia coli* BL21 Rosetta (DE3) (Novagen-Millipore, 70954) and incubated in liquid cultures. After reaching optimum density (0.6 Au at 600nm wavelength) we induced protein construct expression with IPTG, at 0.5 mM final concentration. Cultures were left to further incubate at 20 °C for 16 hrs following IPTG induction. Bacteria were pelleted and re-suspended in lysis buffer (25 mM Tris pH 7.4 • 100 mM NaCl • 2 mM EDTA) additionally supplemented with 0.01% β-mercaptoethanol, and 1 μg/μl lysozyme (final concentrations). Cell integrity was disrupted by sonication using an EpiShear™ Probe Sonicator (in pulses 10 sec ON, 5 sec OFF, 30% amplitude) for 2 min per sample. Protein content was collected as the supernatant of the following centrifugation at 20,000 rpm, 4 °C for 20 min.

Both the GST-bait and His-prey lysates were incubated with Glutathione Sepharose 4B (GE Healthcare/Cytiva, 17-0756-01) for 30 min at 4 °C. Subsequent washes were carried out in High Salt (25 mM Tris pH 7.4 • 500 mM NaCl • 2 mM EDTA) and Low Salt wash buffers (25 mM Tris pH 7.4 • 50 mM NaCl • 2 mM EDTA). After the washes each pre-cleared His-prey lysate was equally distributed to its respective GST bait-enriched beads and samples were co-incubated for 2 hrs at 4 °C. They were subsequently washed with 0.01%-mercaptoethanol-supplemented lysis buffer, and imidazole buffer (lysis buffer recipe + 10 mM imidazole). In preparation for gel loading, samples were finally re-suspended in equal volume of 2x Laemmli solution and denatured at 80 °C for 10 min, prior to SDS-PAGE and subsequent Western Blot. For detection of all His-prey proteins, we used an anti-6xHis antibdy (Abeam, ab18184) at a 1:5000 dilution, while GST-tagged baits were visualised by Ponceau S stain used at a 0.2% working concentration.

### Real-Time qPCR and data analysis

Samples were made from 10 age-matched 2-week-old (after eclosion) adult flies. For the full *Dwg* mutant phenotype (*dwg^8^/dwg^8^), we* used male flies, while heterozygous (*dwg^8^/+*) female flies were used as controls. All procedures were performed according to manufacturer’s guidelines. The RNA was extracted using an Invitrogen PureLink™ RNA Mini Kit (Thermo Fisher, 12183018A). Subsequent steps were performed using 1 μg of total RNA. Genomic DNA was removed by DNAse I digestion (Thermo Fisher, EN0521). For cDNA synthesis we used the RevertAid RT Reverse Transcription Kit (Thermo Fisher, K1691). The qPCR reaction samples were made with the Promega GoTaq qPCR Master Mix (Promega, A6002). The sequences of the forward and reverse primers used in qPCR are given in table below (in 5’ → 3’ direction):

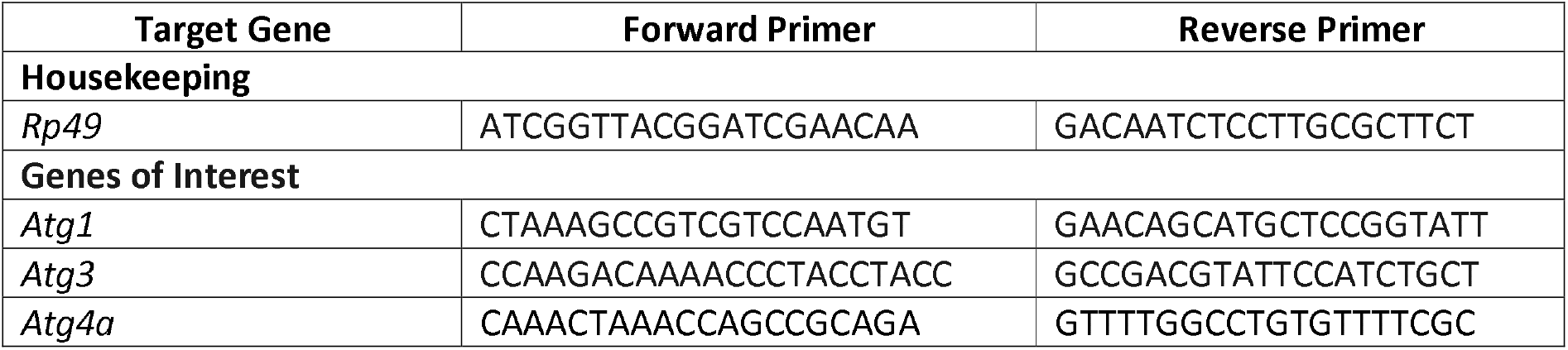
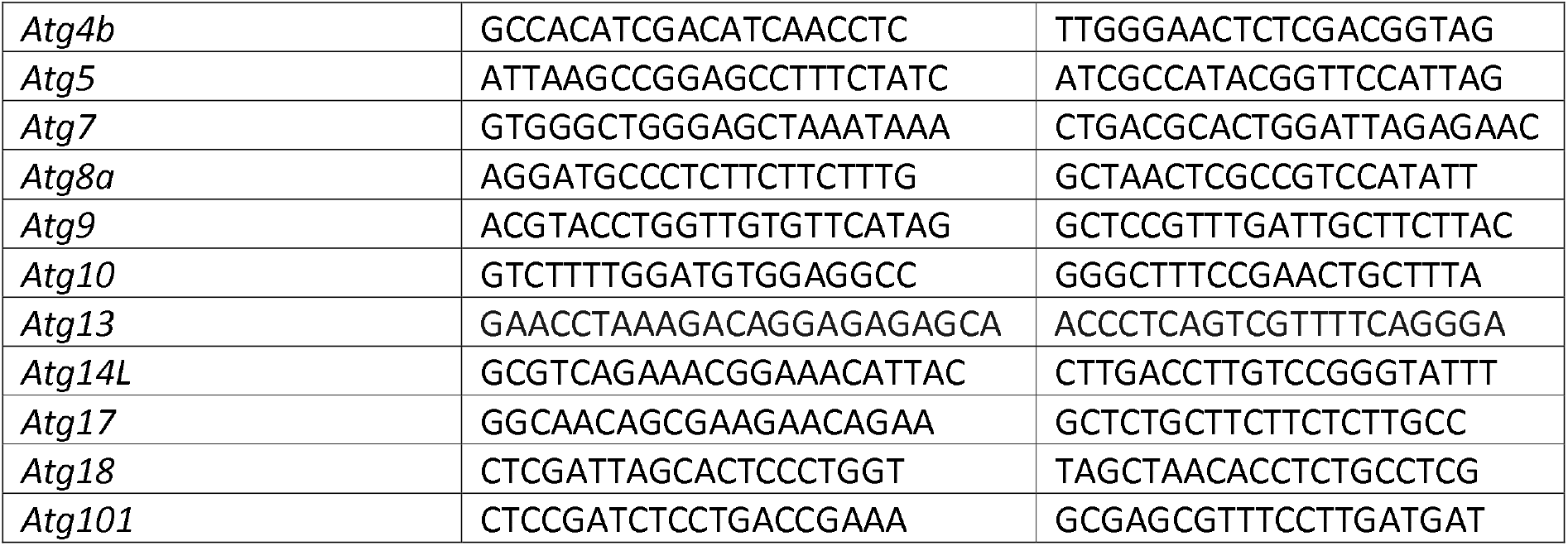
Primer sequences used in qPCR for *Atg* and *Dwg* genes.

Prior to use, we performed a 1:50 working dilution of the cDNA template. Final volume in each well on the qPCR plate per reaction was 25 μl (20 μl primer reaction mix + 5 μl cDNA template). The qPCR assay was performed on a Stratagene Mx3005P (Agilent Technologies) system.

We used the accompanying MxPro software (Agilent Technologies) to obtain the cycle to threshold (Ct) values of the analysed data and employed the 2^ΔΔCt method to calculate gene expression and plot the results in the accompanying graph. Briefly, the Ct of *Rp49* was used as the internal normalizing assay for gene expression within each group, that the Ct of each gene of interest was subtracted from, returning the ΔCt. The genes’ ΔCts of the experimental group were subsequently subtracted from the corresponding ΔCts of the control group giving the ΔΔCt (control group ΔΔCt values consequently set to 0), before being raised to *2^* to give the fold-change in expression of each gene relative to control (control group’s 2^ΔΔCt values consequently set to 1).

### Statistics and figures

All statistical analyses were performed with GraphPad Prism 9 software (version 9.1.0). Figures were assembled in Adobe Photoshop 2020 (version 21.2.5).

**Table 1 legend:** Structural details of experimentally verified LIRCPs in Drosophila using Alpha Fold/Uni-Fold.

## References

Adriaenssens, E., Ferrari, L., & Martens, S. (2022). Orchestration of selective autophagy by cargo receptors. Current Biology: CB, 32(24), R1357–R1371. https://doi.org/10.1016/j.cub.2022.11.002

Evans, R., O’Neill, M., Pritzel, A., Antropova, N., Senior, A., Green, T., Žídek, A., Bates, R., Blackwell, S., Yim, J., Ronneberger, O., Bodenstein, S., Zielinski, M., Bridgland, A., Potapenko, A., Cowie, A., Tunyasuvunakool, K., Jain, R., Clancy, E., … Hassabis, D. (2022). Protein complex prediction with AlphaFold-Multimer (p. 2021.10.04.463034). bioRxiv. https://doi.org/10.1101/2021.10.04.463034

Feng, Y., He, D., Yao, Z., & Klionsky, D. J. (2014). The machinery of macroautophagy. Cell Research, 24(1), Article 1. https://doi.org/10.1038/cr.2013.168

Geng, J., & Klionsky, D. J. (2008). The Atg8 and Atg12 ubiquitin-like conjugation systems in macroautophagy. ‘Protein modifications: Beyond the usual suspects’ review series. EMBO Reports, 9(9), 859–864. https://doi.org/10.1038/embor.2008.163

Han, Z., Zhang, W., Ning, W., Wang, C., Deng, W., Li, Z., Shang, Z., Shen, X., Liu, X., Baba, O., Morita, T., Chen, L., Xue, Y., & Jia, D. (2021). Model-based analysis uncovers mutations altering autophagy selectivity in human cancer. Nature Communications, 12(1), Article 1. https://doi.org/10.1038/s41467-021-23539-5

He, C., & Klionsky, D. J. (2009). Regulation Mechanisms and Signalling Pathways of Autophagy. Annual Review of Genetics, 43(68), 67. https://doi.org/10.1146/annurev-genet-102808-114910.Regulation

Jumper, J., Evans, R., Pritzel, A., Green, T., Figurnov, M., Ronneberger, O., Tunyasuvunakool, K., Bates, R., Žídek, A., Potapenko, A., Bridgland, A., Meyer, C., Kohl, S. A. A., Ballard, A. J., Cowie, A., Romera-Paredes, B., Nikolov, S., Jain, R., Adler, J., … Hassabis, D. (2021). Highly accurate protein structure prediction with AlphaFold. Nature, 596(7873), Article 7873. https://doi.org/10.1038/s41586-021-03819-2

Kalvari, I., Tsompanis, S., Mulakkal, N. C., Osgood, R., Johansen, T., Nezis, I. P., & Promponas, V. J. (2014). ILIR: A web resource for prediction of Atg8-family interacting proteins. Autophagy, 10(5), 913–925. https://doi.org/10.4161/auto.28260

Lamb, C. A., Yoshimori, T., & Tooze, S. A. (2013). The autophagosome: Origins unknown, biogenesis complex. Nature Reviews Molecular Cell Biology, 14(12), 759–774. https://doi.org/10.1038/nrm3696

Li, Z., Liu, X., Chen, W., Shen, F., Bi, H., Ke, G., & Zhang, L. (2022). Uni-Fold: An Open-Source Platform for Developing Protein Folding Models beyond AlphaFold (p. 2022.08.04.502811). bioRxiv. https://doi.org/10.1101/2022.08.04.502811

Ma, Q., Long, S., Gan, Z., Tettamanti, G., Li, K., & Tian, L. (2022). Transcriptional and Post-Transcriptional Regulation of Autophagy. Cells, 11(3), Article 3. https://doi.org/10.3390/cells11030441

Mirdita, M., Schütze, K., Moriwaki, Y., Heo, L., Ovchinnikov, S., & Steinegger, M. (2022). ColabFold: Making protein folding accessible to all. Nature Methods, 19(6), 679–682. https://doi.org/10.1038/s41592-022-01488-1

Nezis, I. P., Lamark, T., Velentzas, A. D., Rusten, T. E., Bjørkøy, G., Johansen, T., Papassideri, I. S., Stravopodis, D. J., Margaritis, L. H., Stenmark, H., & Brech, A. (2009). Cell death during Drosophila melanogaster early oogenesis is mediated through autophagy. Autophagy, 5(3), 298–302.

Pankiv, S., Clausen, T. H., Lamark, T., Brech, A., Bruun, J.-A. A., Outzen, H., Øvervatn, A., Bjørkøy, G., Johansen, T., Øvervatn, A., Bjørkøy, G., & Johansen, T. (2007). P62/SQSTM1 Binds Directly to Atg8/LC3 to Facilitate Degradation of Ubiquitinated Protein Aggregates by Autophagy. Journal of Biological Chemistry, 282(33), 24131–24145. https://doi.org/10.1074/jbc.M702824200

Parzych, K. R., & Klionsky, D. J. (2014). An overview of autophagy: Morphology, mechanism, and regulation. Antioxidants & Redox Signaling, 20(3), 460–473. https://doi.org/10.1089/ars.2013.5371

Rogov, V. V., Stolz, A., Ravichandran, A. C., Rios-Szwed, D. O., Suzuki, H., Kniss, A., Löhr, F., Wakatsuki, S., Dötsch, V., Dikic, I., Dobson, R. C., & McEwan, D. G. (2017). Structural and functional analysis of the GABARAP interaction motif (GIM). EMBO Reports, 18(8), 1382–1396. https://doi.org/10.15252/embr.201643587

Shu, W.-J., Zhao, M.-J., Klionsky, D. J., & Du, H.-N. (2020). Old factors, new players: Transcriptional regulation of autophagy. Autophagy, 16(5), 956–958. https://doi.org/10.1080/15548627.2020.1728611

Yamamoto H, Zhang S, Mizushima N. Autophagy genes in biology and disease. Nat Rev Genet. 2023 Jan 12:1–19.

Xie, Q., Tzfadia, O., Levy, M., Weithorn, E., Peled-Zehavi, H., Van Parys, T., Van de Peer, Y., & Galili, G. (2016). hfAIM: A reliable bioinformatics approach for in silico genome-wide identification of autophagy-associated Atg8-interacting motifs in various organisms. Autophagy, 12(5), 876–887. https://doi.org/10.1080/15548627.2016.1147668

Zhang, Y., & Skolnick, J. (2005). TM-align: A protein structure alignment algorithm based on the TM-score. Nucleic Acids Research, 33(7), 2302–2309. https://doi.org/10.1093/nar/gki524

